# Improved performance of microbial fuel cells through addition of trehalose lipids

**DOI:** 10.1101/339267

**Authors:** Peng Cheng, Rui Shan, Hao-Ran Yuan, Ge Dong, Li-fang Deng, Yong Chen

## Abstract

Electron transfer from microorganisms to the electrode is the key process in microbial fuel cells (MFCs). In this study, a trehalose lipid was added to a *Rhodococcus* pyridinivorans-inoculated MFC to improve the power output by enhancing electron transfer. Upon trehalose lipid addition, the current density and maximum power density were increased by 1.83 times and 5.93 times, respectively. Cyclic voltammetry analysis revealed that the addition of trehalose lipid increased the electron transfer performance, while electrochemical impedance spectroscopy results proved a decrease in internal resistance. Microscopy images showed that the trehalose lipid-treated bacteria interacted more closely with various flagellum-like contacts, while in the pure trehalose lipid (200 mg/L), pores were obviously observed in the cell surface.

**Importance:** Improving the power output of microbial fuel cells by the addition of bio-surfactants have been proved to be a novel method. However, only rhamnolipid and sophorolipid are certified to be effective. Trehalose lipid is a common material in cosmetic and bio-medicine industry. Our research broaden the application of bio-surfactant in MFC and preliminarily explain the mechanism.

**Highlights:**

1. Trehalose lipid enhanced MFC power generation
2. Trehalose lipid decrease MFC internal resistance
3. Pores were observed with the addition of trehalose lipid
4. Addition of bio-surfactant is a promising way to increase MFC performance

## 1. Introduction

Microbial fuel cells (MFCs) are devices that can generate electricity from organic waste, and have drawn significant worldwide research attention (Santoro et al., 2017). Unlike conventional fuel cells, microorganisms act as catalysts on the MFC anode to convert chemical energy to electricity (Akinsemolu, 2018); the bacteria that can generate electricity are called exoelectrogens (Kumar et al., 2015). The bio-process for electricity production in an MFC with an air cathode occurs as follows: (1) the exoelectrogen on the anode oxidizes the substrate to produce electrons and protons, while microbial metabolism occurs in the cytoplasmic matrix; (2) electrons are shuttered to the anode surface from the bacteria; (3) subsequently, electrons gathered at the anode are conducted to the cathode passing through the external load; (4) the electrons and protons are combined with oxygen on the cathode to form water(Zhai et al., 2016, Sun et al., 2016).

The main shortcoming that restricts MFC application is the low power output compared to that of chemical fuel cells. The performance of the MFC significantly depends on the transfer rate from the microbes to the anode. Four mechanisms have been proposed for promoting electron transfer from the exoelectrogen to anode: (1) direct contact with the electrode to convert electrons through cytochrome *(Cyt* c) and membrane bonding proteins (Yang et al., 2014); (2) electrically conductive flagellin-like nanowires (Lovley et al., 2015) (3) the use of electron shutters, a group of electrochemically active substances (Huang et al., 2016), (4) electro-kinesis, by which electrons are transferred to the electrode surface through a rapid wave of fagellin (Lian et al., 2016). Adding exogenous electron transfer mediators (ETMs) is traditionally used to enhance electron transfer (Szoellosi et al., 2015). However, exogenous mediators like ferricyanide and neutral red are expensive, unstable, and sometimes toxic to the microorganisms. Recent studies have found that specific exoelectrogens like *Pseudomonas aeruginosa* and *Geobacter sulfurreducens* can secrete small electrochemically active molecules to transfer electrons (Marsili et al., 2008; Bond et al., 2003). Because bacteria are covered by cell membranes and walls containing non-conductive materials like lipids and peptidoglycan, it is essential to investigate new approaches to the enhance power output by reducing electron resistance.

Recent research have substantiated that bio-surfactants can promote electricity generation in various exoelectrogens (Liu et al., 2012; Song et al., 2015). Moreover, bio-surfactants enhanced electron transfer through cell membranes, remarkably promoting the power output (Wen et al., 2011). The power density and power output were enhanced by 4× and 2.5×, respectively, by the addition of sophorolipid (Shen et al., 2014) and rhamnolipid (Zheng et al., 2015), respectively. However, *Pseudomonas aeruginosa* is the only pure strain confirmed in bio-surfactant application. To broaden the application of bio-surfactants in MFCs, it is necessary to discover additional adaptable exoelectrogens applied in both detected bio-surfactants and effectual biosurfactants.

Bio-surfactants are secondary metabolites of microorganisms with the advantage of good bio-compatibility. Sophorolipids and Rhamnolipids are the most frequently studied surfactants applied in MFCs. Sophorolipids can be synthesized by *Candida bombicola* (Konishi et al., 2008) while rhamnolipids are secreted by *Pseudomonas aeruginosa* (Soberón-Chávez G et al., 2005). Meanwhile, a glycolipid-based bio-surfactant of trehalose lipid with trehalose and carboxylic acid combined as the hydrophilic and hydrophobic groups, respectively, has drawn research attention (Khandelwal et al., 2018). The trehalose lipid is an important bio-surfactant in bio-medicine and cosmetics because of its moisture retention capacity and antibacterial properties (Zaragoza et al., 2013). Trehalose lipids and amber trehalose lipids can be synthesized by *Rhodococcus* bacteria (Philp et al., 2002; Tokumoto et al., 2009). The strain *Rhodococcus pyridinivorans*, which has been proved to be a promising exoelectrogen named HR-1, was separated and cultivated in our lab. The strain is orange-colored with round and smooth colonies, when observed under scanning electron microscopy (SEM) scans, the bacteria are rod-shaped with flagella on the surface. Hence, trehalose lipids were tested in an *R. pyridinivorans-inoculated* MFC to avert bactericidal effects.

Power generation was monitored to study the impact of trehalose lipid addition on the HR-1 inoculated MFC, cyclic voltammetry (CV) was conducted to investigate the electron transfer, electrochemical impedance spectroscopy (EIS) was performed to measure the charge-transfer resistances and the surface morphology of the MFC anode was observed by SEM.

## 2. Materials and methods

### 2.1 MFC setup and operation

A single-chamber air-cathode MFC was assembled from Plexiglas (5 cm ×5 cm × 5 cm, r = 2 cm, 50 mL available volume). Carbon cloth was placed in the anode as the electrode while a membrane cathode, assembled as described by Deng et al., (2016), was loaded with platinum carbon powder dissolved in Nafion (Hesen, Shanghai). Titanium wire was placed between the anode and cathode to conduct electricity. The external resistance was set to 1000 Ω and the MFC was operated at 30 °C in a constant-temperature incubator (Boxun BSC250, China). The logarithmic phase microbe was inoculated into the sterile MFC chamber as anode catalyst. 1 g/L sodium acetate with PBS was the initial anode substrate. As the output voltage dropped below 20 mV after each cycle, the anolyte was replaced with fresh medium. The voltage curves of different experiment groups were obtained after the output was repeatable for three cycles.

### 2.2 Electrochemistry analysis and calculations

The voltage output was monitored by a Keithley multichannel data acquisition instrument (2750, USA), with the electricity current calculated by *I* = *U/R*_ext_. The current and power densities were normalized by the working area of the electrode. To obtain a polarization curve, the external resistance *R*_ext_ was varied from 10 to 10000 Ω by a slide rheostat and the voltage at each resistance was recorded after the output data had stabilized for 2 min.

CV was performed using an electrochemical workstation (CHI1010, Shanghai); the scan rate was 50 mV/s between – 0.8 V and 0.8 *V.* In order to avoid interference, vitamins were removed from the last fed anolyte of the MFC and the chamber was purged with pure filtered N_2_ for 15 min to eliminate O2. During testing, the MFC anode was the working electrode, the MFC cathode was the counter electrode, and an Ag/AgCl (assumed +197 mV vs. standard hydrogen electrode) electrode (MF-2052, BAS) was the reference electrode. All CV tests were conducted at the same temperature and operation conditions (30 °C) and scanning was repeated three times.

### 2.3 SEM observation of exoelectrogen

The bacteria were single-celled soft-structured organisms; it was thus essential to pretreat the sample by the critical point drying method (Xiao et al., 2015). The anode membrane was fixed by 2.5 *%* glutaraldehyde for 4 h, before the sample was washed with 0.1 mol/L phosphate-buffered saline (PBS) for four to six times and then dehydrated by different concentrations of alcohol of 30, 50, 70, 90 and 100%. Finally, the dehydrated anode membrane was replaced by tert-butyl alcohol (TBA).

## 3. Results and discussion

### 3.1 Improved electricity generation by trehalose lipid addition

The voltage data of the single-chamber air-cathode MFC with and without trehalose lipid were recorded. As shown in Fig. 1a, the voltage is significantly enhanced by the addition of trehalose lipid at concentrations below 20 mg/L; the maximum voltage is obtained with the addition of 20 mg/L bio-surfactant. At the trehalose lipid levels of 5, 10 and 20 mg, the voltages are 1.3×, 1.4× and 1.8× higher than those of the control group (0 mg), respectively. However, the voltage curve shows a decrease in the high-concentration experiment group. With the addition of 40 mg/L and 60 mg/L bio-surfactant, the voltage outputs are reduced by 20 mV and 45 mV, respectively. The glycolipid surfactant was similar to the membrane glycoprotein; therefore, according to the principle of similar phase dissolution, the proper addition of trehalose lipid may helped to increase membrane permeability, which resulted in electron transfer enhancement. The adverse effects of trehalose lipid on *R pyridinivorans* MFC were originally expected to act on the bacteria membranes. In other words, the proper concentration of trehalose lipid increased membrane permeability, which enhanced electron transfer, while a high concentration of bio-surfactant harmed the bacteria’s integrality, which reduced the metabolism. Shown in Fig. 1b is a comparison between the untreated MFC and 20 mg/L-treated MFC regarding voltage and power densities. At the same fixed external resistance of 1000 Ω, with the addition of optimal bio-surfactant, the MFC shows the maximum power density of 0.25 mW/cm^2^, which is 3.3× that of the untreated MFC (0.075 mW/cm^2^). Meanwhile, bio-surfactants can reduce surface tension and retain moisture; these features are confirmed by the treated MFC reaching its maximum voltage earlier than the control, which possibly indicates microorganism biofilm formation on the anode in the initial phase.

**Figure 1.**
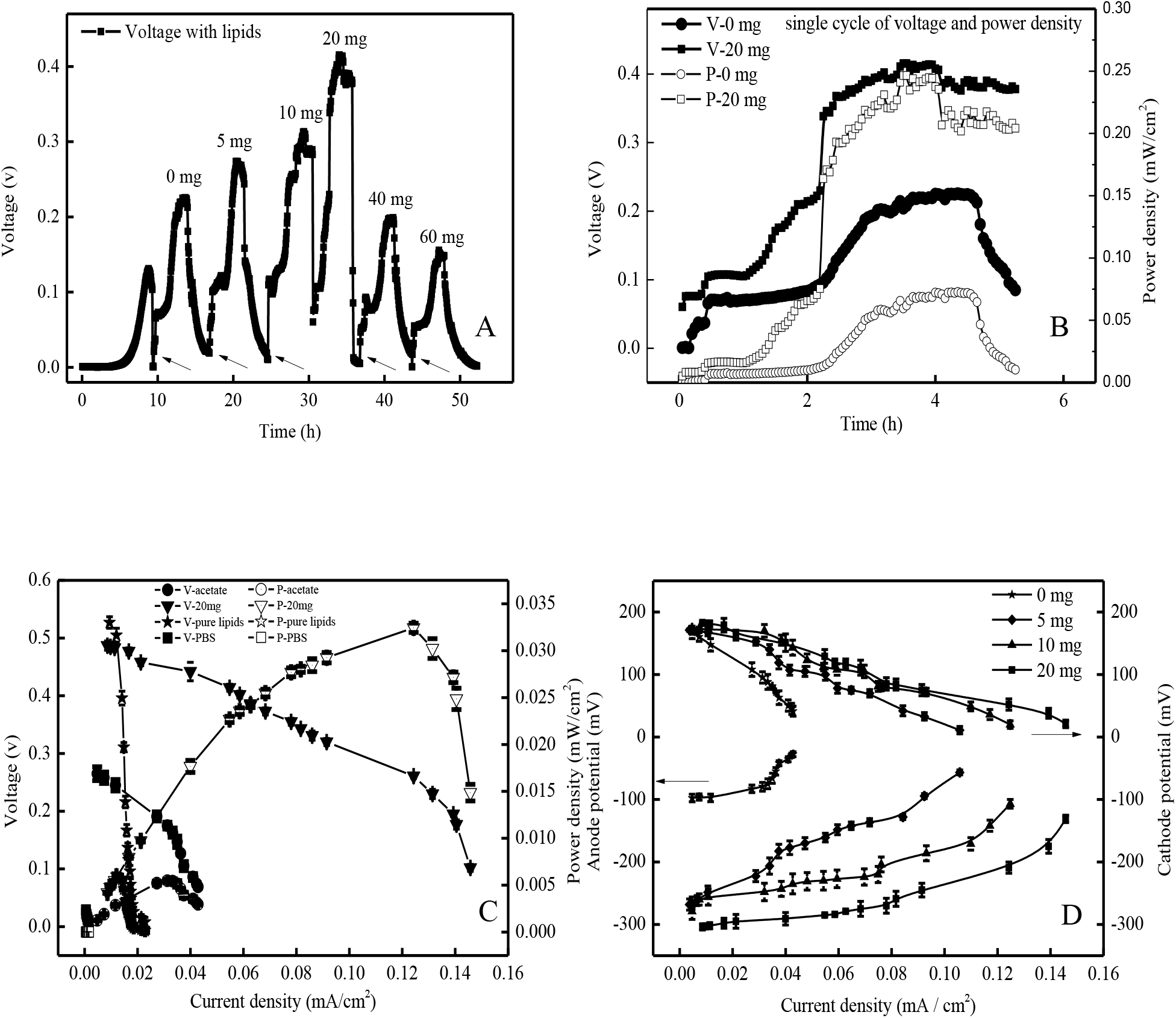
A. Voltage output of the *R. pyridinivorans* sp. strain HR-1-inoculated MFC under different concentrations of trehalose lipid of 0, 5, 10, 20, 40, and 60 mg/L; B. Voltage and power density comparison between untreated and 20 mg/L bio-surfactant-treated MFCs; C. Polarization curves of the MFC with only 1 g/L acetate, 1 g/L acetate with 20 mg/L trehalose lipid, pure trehalose lipid anolyte, and pure PBS.; D. Electrode potential of the MFC under trehalose lipid addition at concentrations of 0, 5, 10, and 20 mg/L.

Polarization curves and power density measurements obtained by varying the external resistance were used to investigate the effect of trehalose lipid on the electricity generation in the MFC. Shown in Fig. 1c are the polarization curves of 1 g/L acetate, pure PBS, 1 g/L acetate with 20 mg/L bio-surfactant, and pure bio-surfactant. With the addition of pure PBS or trehalose lipid, the MFC can generate only negligible electricity. The results also show that neither PBS solution nor trehalose lipid is an electron donor for electricity generation. However, with trehalose lipid addition, the power density shows a much higher polarization in the MFC, indicating better energy conversion efficiency compared to that of the control (Srikanth et al., 2012). In the control group without bio-surfactant, the MFC shows the maximum power density of 0.0049 mW/cm2 at the current density of 0.035 mA/cm^2^; with the addition of 20 mg/L trehalose lipid, themaximum power density of 0.033 mW/cm^2^ is obtained at the current density of 0.128 mA/cm^2^, which is 6.7× higher than of the control. Meanwhile, the maximum current density is increased from 0.044 mA/cm^2^ to 0.148 mA/cm^2^, indicating a decrease in electron transfer resistance under the effect of the bio-surfactant (Yong et al., 2013). According to Ohm’s law, the internal resistance can be calculated by the linear area of the polarization curve. The estimated internal resistance with the trehalose lipid addition is 366 Ω, which is only 43 *%* of that of the untreated MFC (852 Ω). The results reveal that, through the addition of trehalose lipid, the charge-transfer resistance is significantly reduced, suggesting that the power output and energy conversion efficiency are enhanced by the bio-surfactant addition.

The electrode potential was measured to evaluate the effect of trehalose lipid on the anode and cathode of the MFC. Shown in Fig. 1d are the electrode potentials with the current density of the MFC operated under different concentrations of trehalose lipid addition (0, 5, 10, and 20 mg/L). With bio-surfactant addition, both the anode and cathode potentials are obviously increased compared to those of the control group. The bio-surfactant can reduce surface resistance, which benefits oxygen infusion and bacteria growth (Wen et al. 2010). Moreover, at the current density of 0. 059 mA/cm^2^, the anode potentials are 149, 227, and 284 mV and the cathode potentials are 88, 108, and 118 mV with trehalose additions of 5, 10, and 20 mg/L, respectively. However, as the concentration of trehalose lipid is increased, the cathode potential does not change significantly compared to the anode potential with the increase of the current density. The effect of the bio-surfactant on the electrode potential promoted the power output of the MFC. Moreover, the results of the polarization curves and anode potential indicate that the anode performance under trehalose lipid addition is mainly responsible for the overall power generation.

### 3.2 Electrochemical performance with trehalose lipid

CV was performed to examine the electrochemically active substances during electron transfer at a stable MFC output voltage (Mohanakrishna et al., 2018). Shown in Fig. 2a are the CV curves of MFCs with different concentrations of trehalose lipid. It is obvious that the redox pair peaks are centered at −0.36 V and −0.28 V The peak currents are enhanced as the addition is increased, indicating that the trehalose lipid can promote the electrochemical activity of the MFC. At the addition of 20 mg/L bio-surfactant, the peak current is 0.93 mA, three times higher than that of the initial MFC. Fig. 2b and Fig. 2c show comparisons between 1 g/L acetate-fed MFC and the pure trehalose lipid addition and between pure PBS anolyte with and without bio-surfactant addition, respectively. It is obvious that no distinct peaks are formed during the CV scans in the pure bio-surfactant group and in the pure PBS anolyte without trehalose lipid addition, which implies that the trehalose lipid has no electrochemical activity and cannot serve as an electron donor for the MFC. This result excludes the possibility of acting on the bacteria metabolism by impacting the chemical oxygen demand (COD) and illustrates that the addition of trehalose lipid to the MFC positively affects electron transfer on the anode.

**Figure 2.**
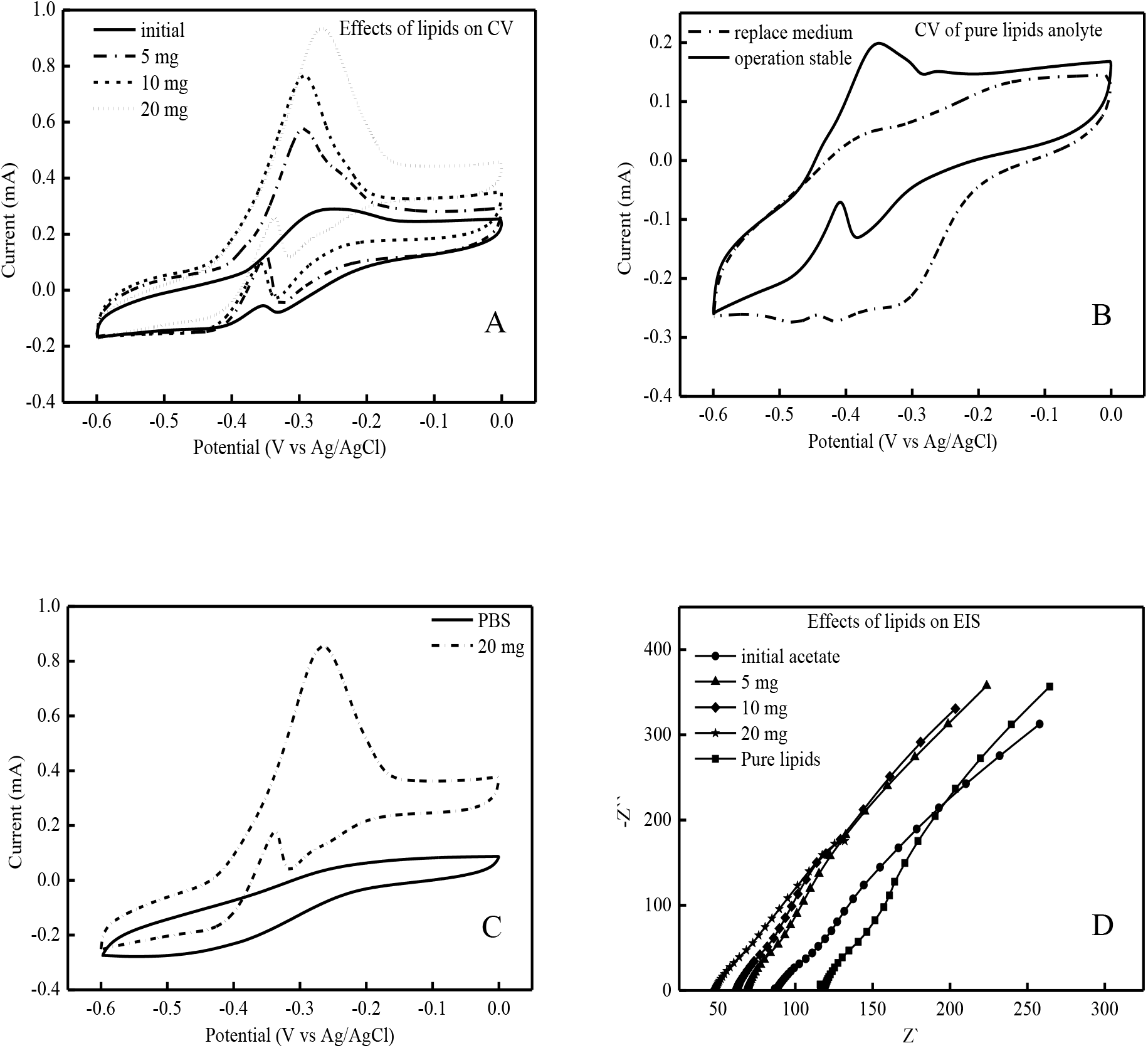
A. CV analysis of MFC at stable output voltage under different concentrations of trehalose lipid addition. The bio-surfactant concentrations are 0, 5, 10, and 20 mg/L; B. CV analysis of pure trehalose lipid anolyte on turnover phase; C. CV peak comparison of 20 mg/L trehalose lipid-treated and untreated MFCs; D. EIS analysis of the MFC anode with different substrates.

To further investigate the impact of the bio-surfactant on the MFC, EIS was performed to measure the charge-transfer resistance. Fig. 2d shows the impedance spectra for anodes with different substrates. Unlike the experimental group with acetate, that with pure trehalose lipid has a relatively high resistance. The CV results showed that the trehalose lipid did not act as an electron donor, which decreased the electron concentration and thus increased the internal resistance. Nevertheless, the experimental groups with different concentrations of bio-surfactant show decreases in the internal charge transfer; the variation tendencies agree with the CV results. Based on the Nyquist plots of the EIS curves, the internal resistance of the MFC with 20 mg/L trehalose lipid addition is only 56% that of the untreated MFC. The charge-transfer resistance analysis was similar to the polarization linear calculation, verifying the conjecture that the trehalose lipid affected bacteria at the anode to enhance electron transfer, which promoted the power output.

### 3.3 Effect of trehalose lipid on bacteria surface

Previous studies have illustrated that through surfactant addition, anodic bacteria attachment was enhanced (Zhang et al., 2017). However, bacteria were perforated when treated with cationic reagents (Liu et al., 2012). In this experiment, the MFC power output was promoted when the MFC bacteria were treated with trehalose lipid at the concentration of 20 mg/L, while the output was restricted at higher-concentration additions of the bio-surfactant. SEM observation was conducted to determine the adverse results.

Fig.3a shows the surface morphology of the carbon cloth before inoculation; the carbon cloth is smooth without any impurities. When treated with the electron donor and inoculated with *R. pyridinivorans* sp. strain HR-1, the successfully initiated MFC anode is shown to bear numerous bacteria (Fig. 3c). The bacteria are rod-shaped with smooth surfaces and gathered on the anode surface (Fig. 3f). In contrast, the surface of the MFC anode with 20 mg/L trehalose lipid is more diversified; the bacteria are more plump and interconnected by flagella-like substances (Fig. 3b and Fig. 3e). The microorganism increases in volume if the inner osmotic pressure is decreased; the osmotic pressure decrease may be caused by pore formation upon bio-surfactant addition (Yates et al., 2012). In the meantime, the trehalose lipid surfactant lowers the surface tension and enhances contact between bacteria. The pore formation conjecture is proved by Fig. 3d, where obvious pores are observed on the bacteria surfaces. The surface morphology with pure trehalose lipid addition is more like the anode with attached bacteria and no bio-surfactant addition, which agrees with the power generation results and electrochemical analysis. The plumped bacteria and their surface connections both promote electron transfer, while the pure trehalose lipid-treated MFC and untreated MFC have higher internal resistances. In conclusion, pores formed by the bio-surfactant can act as shutter channels to enhance electron transfer from the cytoplasm to the extracellular domain, but high surfactant concentrations can break the cell structure, which can lead to metabolism weakening.

**Figure 3.**
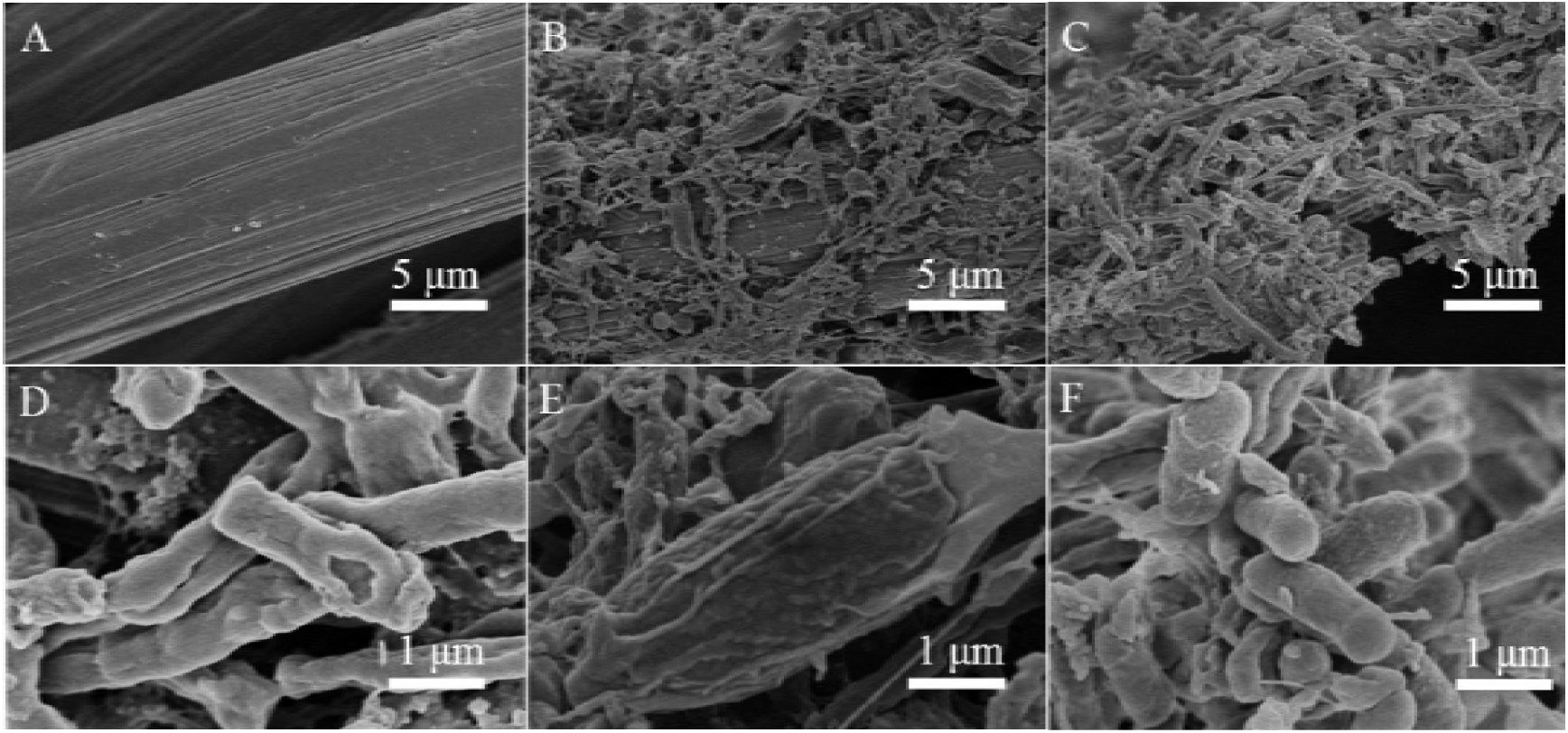
Surface morphology of anode under different treatments. *A.* Blank carbon cloth; B. *R. pyridinivorans* sp. strain HR-1-inoculated MFC anode with 1 g/L acetate and 20 mg/L trehalose lipid; C. Untreated MFC anode surface; D. Pure trehalose lipid anolyte-treated MFC anode; E. *R. pyridinivorans* sp. strain HR-1 inoculated MFC anode with 1 g/L acetate and 20 mg/L trehalose lipid; F. Untreated MFC anode surface.

## Conclusion

In this study, the addition of a bio-surfactant (trehalose lipid) for improving power production and reducing the resistance in a *Rhodococcus pyridinivorans* sp. strain HR-1 MFC was successful performed. With the addition of 20 mg/L trehalose lipid, the maximum power density of 1 g/L acetate-fed MFC was increased from 0.05 mW/cm^2^ to 0.3 mW/cm^2^ (a six-fold enhancement). Pores were observed on the bacteria surfaces and resulted in improvements in the open-circuit voltage and power density of the MFC. This study confirmed that the addition of bio-surfactants to MFCs could enhance bioelectricity generation.

## Acknowledgements

We gratefully acknowledge financial support from the National Natural Science Foundation of China (51406207 and 51676194), Scientific Research Equipment Development Project of Chinese Academy of Sciences (YZ201516), Science and Technology Project of Guangdong Province, China (2016201604030077), and the Science and Technology Project of Guangdong Province, China (2016A010105017).

